# Combination of deep neural network with attention mechanism enhances the explainability of protein contact prediction

**DOI:** 10.1101/2020.09.04.283937

**Authors:** Chen Chen, Tianqi Wu, Zhiye Guo, Jianlin Cheng

## Abstract

Deep learning has emerged as a revolutionary technology for protein residue-residue contact prediction since the 2012 CASP10 competition. Considerable advancements in the predictive power of the deep learning-based contact predictions have been achieved since then. However, little effort has been put into interpreting the black-box deep learning methods. Algorithms that can interpret the relationship between predicted contact maps and the internal mechanism of the deep learning architectures are needed to explore the essential components of contact inference and improve their explainability. In this study, we present an attention-based convolutional neural network for protein contact prediction, which consists of two attention mechanism-based modules: sequence attention and regional attention. Our benchmark results on the CASP13 free-modeling (FM) targets demonstrate that the two attention modules added on top of existing typical deep learning models exhibit a complementary effect that contributes to predictive improvements. More importantly, the inclusion of the attention mechanism provides interpretable patterns that contain useful insights into the key fold-determining residues in proteins. We expect the attention-based model can provide a reliable and practically interpretable technique that helps break the current bottlenecks in explaining deep neural networks for contact prediction.

## 1. Introduction

Prediction of residue-residue contacts in proteins plays a vital role in the computational reconstruction of protein tertiary structure. Recently, advancements in the mathematical and statistical techniques for inter-residue coevolutionary analysis provide essential insights for correlated mutation-based contact prediction, which is now becoming a critical component to generate input features for machine learning contact prediction algorithms. For instance, in the recent 13th Community-Wide Experiment on the Critical Assessment of Techniques for Protein Structure Prediction (CASP13) contact prediction challenge, significant improvements have been achieved due to the integration of both inter-residue coevolutionary analysis and novel deep learning architectures ^1–4^.

A variety of deep learning-based models have been proposed to improve the accuracy of protein contact prediction since deep learning was applied to the problem in 2012 CASP10 experiment^5^. Many of these methods leverage the contact signals derived from the direct coupling analysis (DCA). Most DCA algorithms ^6–9^ generate correlated mutation information between residues from multiple sequence alignments (MSAs), which is utilized by the deep convolutional neural networks in the format of 2D input feature maps. For example, RaptorX-Contact^10^, DNCON2^11^, and MetaPSICOV^12^ are a few early methods that apply the deep neural network architectures with one or more DCA approaches for contact prediction. The connection between the two techniques underscores the importance of explaining the contribution of patterns in coevolutionary-based features to the deep learning-based predictors.

Despite the great success of deep learning-based models in a variety of tasks, this approach is often treated as black-box function approximators that generate classification results from input features. Since the number of parameters in a network is somewhat proportional to its depth, it is infeasible to extract human-understandable justifications from the inner mechanisms of deep learning without proper strategies. Saliency maps and feature importance scores are widely used approaches for model interpretation in machine learning. However, due to the unique characteristic of contact prediction, these methods involve additional analysis procedures that require far more computational resources than other typical applications. For example, the saliency map for a protein with length L requires L×L times of deconvolution operations in a trained convolutional neural network since the output dimension of contact prediction is always the same as its input. While this number can be reduced by choosing only positive labels for analysis, the whole saliency map is still much harder to determine since the many DCA features fed to the network have higher dimensions than the traditional image data. For example, RaptorX-Contact ^10^, one of the state-of-the-art contact predictors, takes 2D input with a size of L×L×153. The recent contact/distance predictor DeepDist^13^ takes input with size up to L×L×484. The very large input size for contact prediction makes it difficult to use these model interpretation techniques.

Recently, the attention mechanism has been applied in natural language processing (NLP)^14,15^, image recognition^16^, and bioinformatics ^17,18^. The attention mechanism assigns different importance scores to individual positions in its intermediate layer so that the model can focus on the most relevant information within the input. In 2D image analysis, the attention weights for individual positions on the contact map allow the visualization of critical regions that contribute to the final predictions. In addition, these weights are generated during the inference step, without requiring additional computation procedures after the prediction of a contact map. Hence, the attention mechanism is a suitable technique to facilitate the interpretation of protein contact prediction models.

In this article, we propose an attention-equipped deep learning method for protein contact prediction, which adopts two different architectures of the attention targeted for interpreting 2D and 1D input features, respectively. The regional attention utilizes the n×n region around each position of its input 2D map while the sequence attention utilizes the whole range of its 1D input. The regional attention module is implemented with a specially designed 3D convolutional layer so that training and prediction on large datasets can be performed with high efficiency. The sequence attention is similar to the multi-headed attention mechanism applied in the NLP tasks. We show that our attention-based methods can achieve reliable accuracy while improving the interpretability of the model and providing potentially useful insights for the identification of residues that are critical in determining protein folds.

## 2 Materials and Methods

### 2.1 Overview

The overall workflow of this study is shown in **Figure 1**. We use the combined predictions from two neural network modules of different attention mechanisms (sequence attention and regional attention) to predict the contact map for a protein target. Both modules take two types of features as inputs: the pseudolikelihood maximization matrix (PLM) ^8^ generated from multiple sequence alignment as a coevolution-based 2D feature and the position-specific scoring matrix (PSSM) which provides the representation of the sequence profile for the input protein sequence. The outputs of the two modules are both L X L contact maps with scores ranging from 0 to 1, where L represents the length of the target protein. The final prediction is produced by the ensemble of two attention modules. We implemented our model with Keras (https://keras.io). For the evaluation of the predicted contacted contact map, we primarily focus on long-range contacts (sequence separation between two residues: n ≥24).

**Figure 1.**
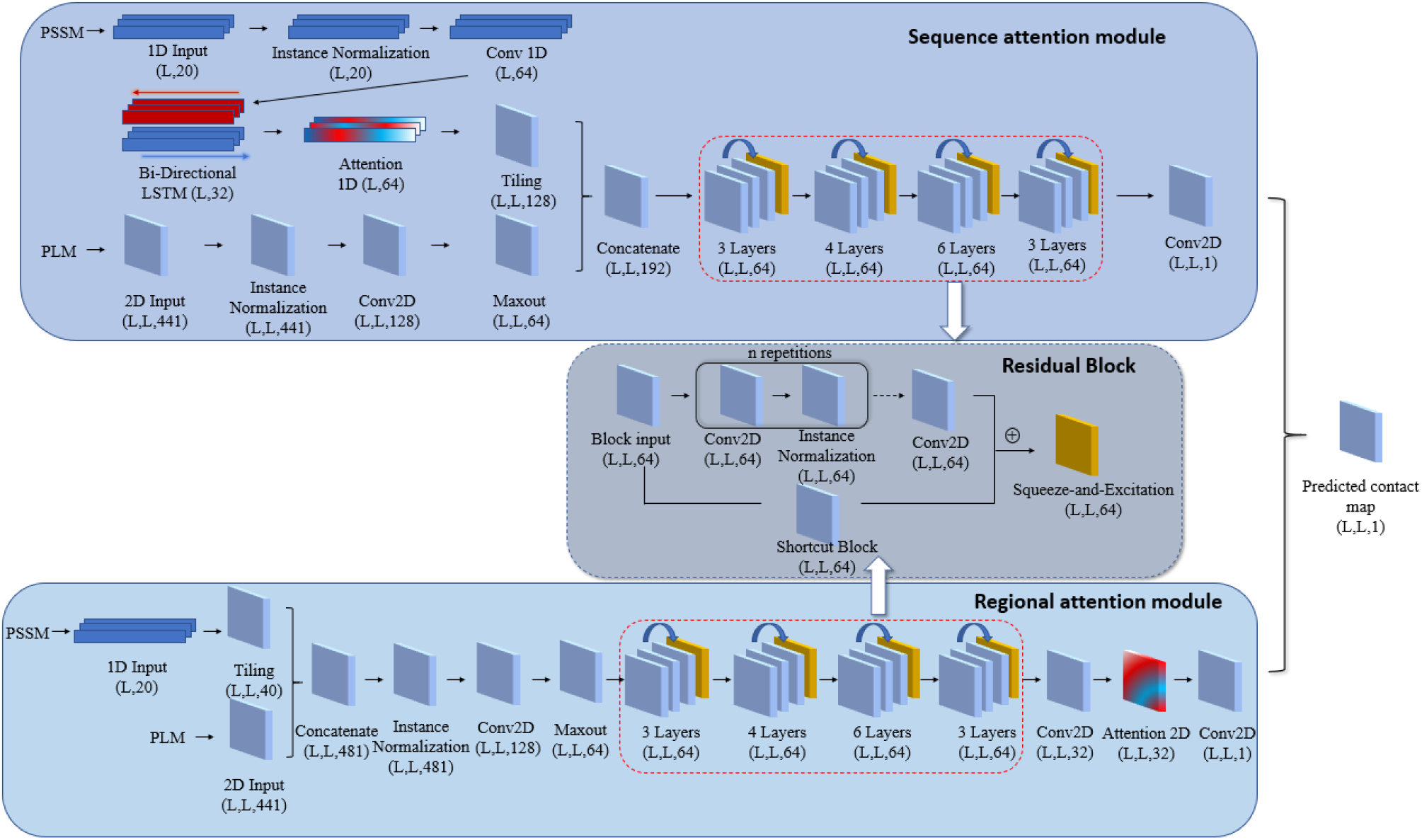
An overview of the proposed attention mechanism protein contact predictor framework. The architecture of the deep neural network employed with two attention modules: In the sequence attention module, the 1D input (PSSM) first goes through the 1D convolution network and bidirectional long- and short-term memory network (LSTM). Then the attention mechanism is applied to the LSTM output. The 2D input (PLM) is first processed with the 2D convolutional neural network and the Maxout layer. The 1D input is then tiled to 2D format so that it can be combined with the 2D input. The concatenated inputs then go through a residual network with 4 residual blocks consist of 3, 4, 6, 3 repetitions of 2D convolution layers, respectively. In regional attention networks, the 1D inputs are first tiled to 2D format and concatenated with the 2D input. The combined inputs are then processed similarly with the sequence attention module, except for the additional 2D attention layer before the last convolution layer. The average of the outputs from the two modules is used as the final predicted contact map.

### 2.2 Datasets

We select targets from the training protein list used in DMPfold ^19^ and extract their true structures from the Protein Data Bank (PDB) to create a training dataset. After removing the redundant proteins that may have >25% sequence identity with any protein in the validation dataset and test dataset, 6463 targets are left in the training dataset. The validation set contains 144 proteins used to validate DNCON2 ^11^. The blind test dataset is built from 31 CASP13 free modeling (FM) domains. The CASP13 test dataset contains new proteins that have no sequence similarity with both the training and test datasets at all.

### 2.3 Input Feature Generation

For each protein sequence, we use two features as inputs for the deep learning model: PLM and PSSM. The PLM is generated from multiple sequence alignments (MSAs) produced by DeepMSA [16]. The sequence databases used in the DeepMSA homologous sequences search include Uniclust30 (2017-10) ^20^, Uniref90 (2018-04) and Metaclust50 (2018-01) ^21^, our in-house customized database which combines Uniref100 (2018-04) and metagenomics sequence databases (2018-04), and NR90 database (2016). All of the sequence databases used for feature generation were constructed before the CASP13 experiment (e.g. before the CASP13 test dataset was created). DeepMSA combines iteratively homologous sequence search of HHblits^22^ and Jackhmmer^23^ on the sequence databases to compute MSAs for feature generation. It performs trimming on the sequence hits from Jackhmmer with a sequence clustering strategy, which reduces the search time of the HHblits database construction for the next round of search.

### 2.4 Deep Network Architectures

Our model consists of two primary components, the regional attention module, and the sequence attention module (**Figure 1**). The two modules include the attention layers, normalization layers, convolution layers and residual blocks. The outputs of these two modules are combined to generate the final prediction. Below are the detailed descriptions of each module with an emphasis on the attention layers.

#### 2.4.1 Sequence attention module

In the sequence attention module (**Figure 1**), the 1D PSSM feature first goes through an instance normalization layer ^24^ and a 1D convolution operation, which is followed by a Bi-Directional LSTM layer in which the LSTM operations are applied on both forward and reverse directions of the inputs. The output vectors on both directions are concatenated. The outputs are then fed into a multi-headed scaled dot product attention layer (**Figure 2a**). Three vectors required for the attention mechanism: Q(Query), K(Key), and V(Value) are generated from different linear transformations of the input of the attention layer. The attention output *Z* is computed as:

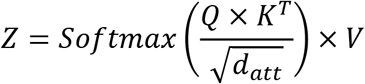

where *d*_*att*_ represents the dimensions of Q, K and V. The attention 1D operation assigns different weights to the transformed 1D feature so that the critical input region for the successful prediction can be spotted. The attention output Z is then tiled to 2D format by the repetition of columns and rows for each dimension.

**Figure 2.**
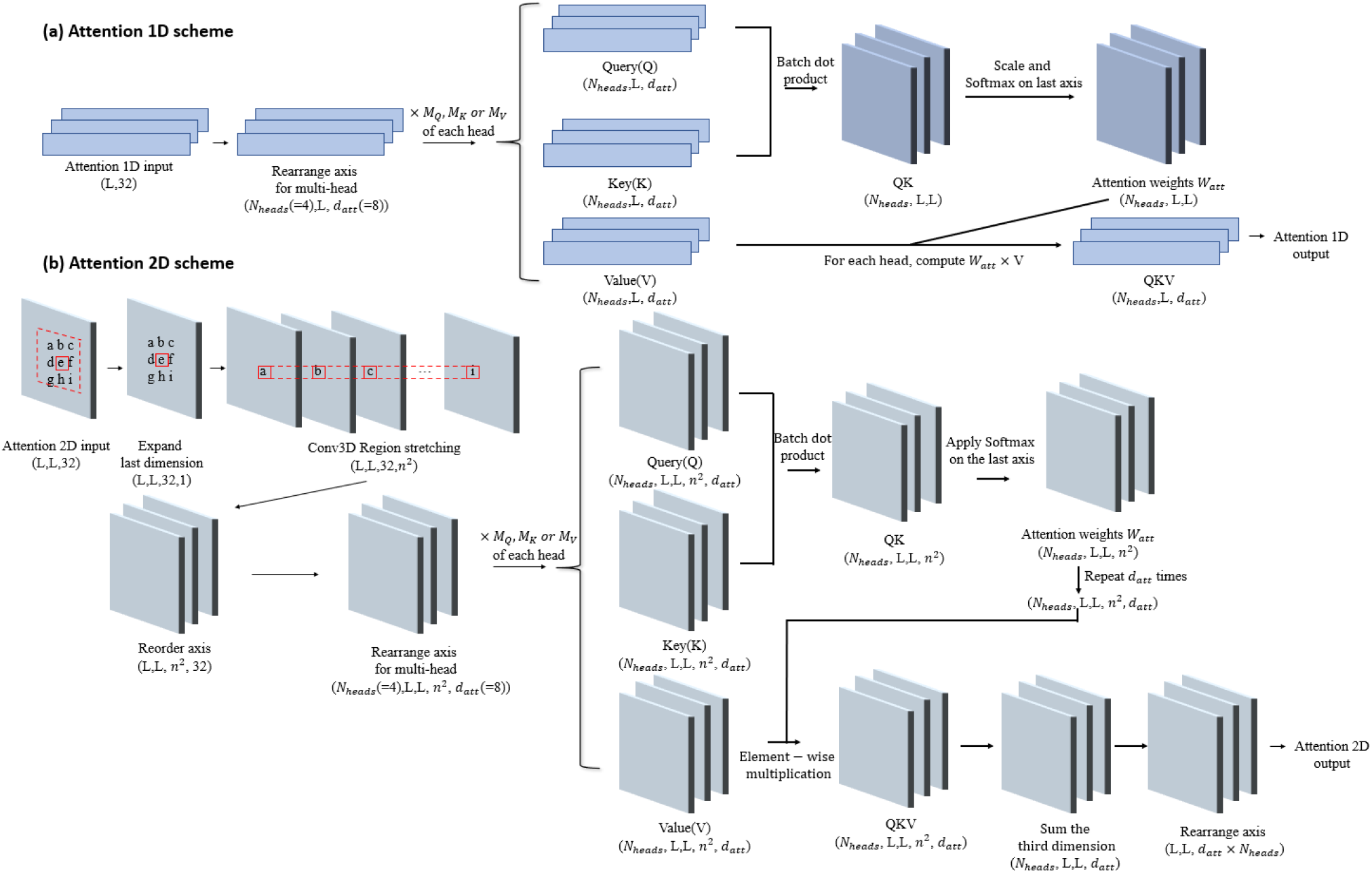
Schematic illustration of 1D and 2D attention mechanism. **(a)** The scheme for 1D attention mechanism. The input is first transformed into a vector of size (*N*_*heads*_, L, *d*_*att*_) for the efficient multi-headed attention implementation. For each head, the vector of size (L, *d*_*att*_) is multiplied to three different trainable matrices of size (*d*_*att*_, *d*_*att*_) to generate Query(Q), Key(K), and Value(V). Different heads have their own transformation matrices for Q, K and V. Q and K first go through a batch dot product operation, resulting in a new vector QK with size (*N*_*heads*_, L, L). QK is then scaled and normalized with Softmax function on the last axis, which becomes the attention score *W*_*att*_. *W*_*att*_ × *V* for each head becomes the 1D attention output. **(b)** The scheme for 2D attention mechanism. The 2D input is first transformed with a 3D convolution and becomes a stretched vector of size (L, L, 32, *n*^2^). It is then computed with the similar attention operation as the 1D attention scheme on the last axis. In practice, different heads for both sequence and regional attention are treated as samples in a batch to improve training speed.

The 2D feature PLM first goes into the instance normalization and a ReLU activation ^25^. It is then processed by a convolutional layer with 128 kernels of size 1×1 and a Maxout layer ^26^ to reduce the input dimension from 128 to 64. The 2D inputs are concatenated with the tiled attention output and go into the residual network component. The final output of the sequence attention module is generated from a 2D convolution layer with a filter of size (1,1) and Sigmoid activation, resulting in output of size L×L.

#### 2.4.2 Regional attention module

The regional attention module (**Figure 1**) takes inputs from the PLM matrix and the tiled 2D PSSM feature. The two features are concatenated at the beginning of the module and are processed in the same way as the 2D PLM input of the sequence attention module. The residual network component with the same configuration (described in 2.4.3) as in the sequence attention module is also applied. The last residual block is followed by a convolutional layer with 32 filters, and the results are used as the input of the attention 2D layer.

The input shape of the attention 2D layer (**Figure 2b**) is (L, L, 32). It is converted by a 3D convolution layer (Region Stretching layer) with specially designed filters so that the output has shape (L, L, 32, *n*^2^), where *n* is the dimension of the attention region for each position in the 2D input. The purpose of this layer is to make the last dimension of its output represent the flattened *n*×*n* region around each element of the original input (in our model *n* is set to 5). The Region Stretching layer has *n*^2^ filters with shape *n*×*n*. For the *i*-th filter of the layer, the weight of the *i*-th element (flatten in row-major order) in the *n*×*n* area is always set to 1 with all other positions set to 0. We repeat these filters 32 times so that the stretching operation is applied to all dimensions of the input. The weights of these filters will not be changed during training. This operation can leverage the highly optimized convolution implementation in Keras and is much more efficient than the explicit implementation. The corresponding Q, K, and V vectors for the attention mechanism are computed from the transformed output of 3D convolution. The scaling and Softmax normalization are applied to the last dimension for the products of Q and K so that different attention weights can be assigned to the *n*×*n* surrounding area for each position on the L×L map. As a result, the output of each position on the feature map will be a weighted sum of their surrounding regions. After the 2D attention layer, the output of the regional attention module is generated from a 2D convolutional layer with a filter of size (1,1) and the Sigmoid activation.

#### 2.4.3 Residual network architecture

Both attention modules have the same 34-layer residual network architecture consisting of four residual blocks differing in the number of internal layers (**Figure 1**). Each residual block is composed of several consecutive instance normalization layers and convolutional layers with 64 kernels of size 3×3. The final values of the last convolutional layer are added to the output of a shortcut block, which is a convolutional layer with 64 kernels of size 1×1. A squeeze-and-excitation (SE) block^27^ is added at the end of each residual block. The SE operation weights each of its channels differently by a trainable 2-layer dense network when creating the output feature maps, so that channel-wise feature responses can be adaptively recalibrated.

### 2.5 Training

The training of the deep network is performed with the customized Keras data generators to reduce the memory requirement. The batch size is set to 1 due to the large size of feature data produced from long protein sequences. A normal initializer ^28^ is used to initialize the weights of the layers in the network. Adam optimizer^29^ is used for training, with the initial learning rate set to 0.001. For epochs ≥ 30, the optimizer is switched to stochastic gradient descent (SGD)^30^, with a learning rate of 0.01 and a momentum of 0.9. At the end of each epoch, the current weights are saved, and the precision of top L/2 long-range contact predictions (e.g. predicted contacts with sequence separation >= 24) on the validation dataset is evaluated. The training process is terminated at epoch 60, and the epoch with the best performance on the validation dataset is chosen for the final blind test.

## 3 Results

### 3.1 The contact prediction accuracy on the CASP13 dataset

We evaluate the performance of our models on 31 CASP13 free-modeling (FM) targets. According to the definition from CASP13, a pair of residues are considered to be in contact if the distance between their C_β_ atoms in the native structure is less than 8.0 Å. By convention, long-range contacts are defined as contact pairs in which the sequence separations between the two residues of the contacts are larger than or equal to 24 residues. The sequence separation for medium-range is between 12 and 23 and short-range between 6 and 11 residues. Following a common standard in the field ^1^, we evaluate the precision of top L/n (n = 1, 2, 5) predicted long-range contacts. In addition to evaluating the overall performance of the combined model, we benchmarked the predictions from the two independent attention modules. The evaluation results are shown in **Table 1**.

**Table 1.** Precision (%) of the top L/5, L/2 and L predicted long-range contacts on the CASP13 dataset.

| Model | Short-range |  |  | Medium-range |  |  | Long-range |  |  |
| --- | --- | --- | --- | --- | --- | --- | --- | --- | --- |
|  | Top-L/5 | Top-L/2 | Top-L | Top-L/5 | Top-L/2 | Top-L | Top-L/5 | Top-L/2 | Top-L |
| Sequence attention | 58.26 | 41.51 | 27.95 | 63.10 | 45.87 | 32.51 | 64.46 | 52.13 | 39.82 |
| Regional attention | 61.08 | 41.95 | 28.38 | 65.46 | 48.00 | 33.65 | 67.32 | 54.15 | 40.96 |
| Combined model | 60.94 | 42.69 | 28.83 | 66.45 | 48.45 | 34.19 | 70.73 | 55.88 | 42.64 |
| Baseline model | 59.00 | 42.73 | 28.17 | 64.29 | 48.01 | 33.05 | 66.31 | 49.42 | 36.40 |

The combined model outperforms each individual attention model and model without attention mechanism for top L/5, top L/2, and top L predicted contacts in medium and long-range. For instance, the top L/5 long-range precision of the combined model is 70.73%, higher than both the sequence attention module (64.46%) and the regional attention module (67.32%) as well as the baseline model that without either of the attention mechanisms.

We also find that the predictive improvements in combining the two attention modules are from the predictions with high confidence scores. **Figure 3a** and **3b** illustrate the receiver operating curve (ROC) and Precision-Recall curves (PR curve) of the three models on targets for evaluation. The area under the curve (AUC) for ROC curve and PR curve of all three models has similar trends. **Figure 3c** and **3d** show the ROC and PR curves of the union of residue pairs from top-L/5 scores in any of the three models. For AUC of both curves, the combined results have a higher score (0.7888 for ROC curve and 0.8031 for PR curve) than the sequence attention model (0.7614 for ROC curve and 0.7935 for PR curve) and the regional attention model (0.7769 for ROC curve and 0.7907 for PR curve). The improved result of combining the two attention models indicates that they are complementary.

**Figure 3.**
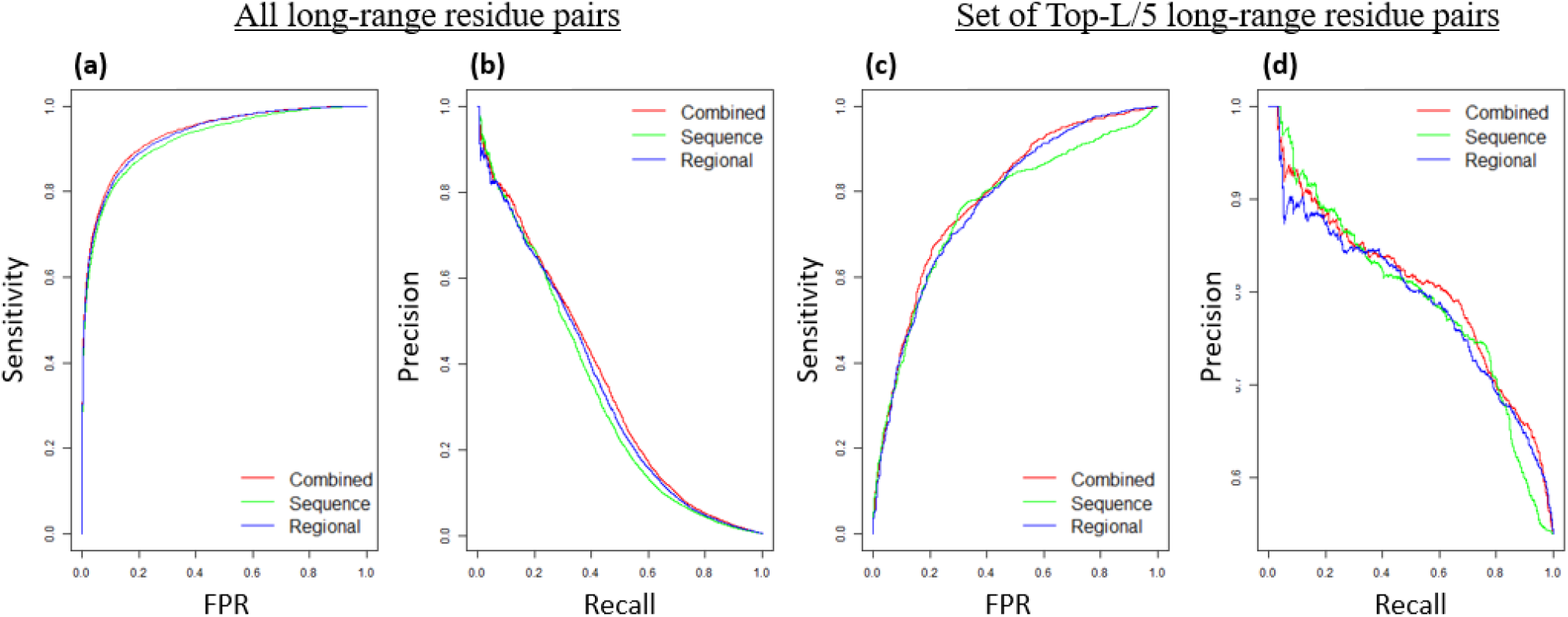
Prediction performance curves of the sequence attention model, regional attention model, and combined model. **(a)** ROC curve for all long-range contact predictions. **(b)** Precision-Recall curve for all long-range contact predictions. **(c)** ROC curve for all residue pairs that appear in the union of residue pairs from top-L/5 scores in any of the three models. **(d)** Precision-Recall curve for all residue pairs that appear in the top-L/5 scores in any of the three models.

### 3.2 Comparison of the predictive performance of two attention modules

We compare the performance of the two attention modules for each target in **Figure 4**. The results show that the precision scores of the two attention modules have a strong correlation (Pearson Correlation Coefficient=0.78) among all targets. As expected, most of the targets with high prediction precision in the combined model are those with high precision scores in both attention modules. Interestingly, there are cases in which the combined predictions acquire an improved performance when the two attention modules perform very differently. For example, the top-L/5 precision score of T1008_D1 reaches 93.33% in the combined model, higher than the sequence module (46.67%) and the regional module (80.00%). Similarly, the top-L/5 precision score of T0957s2 reaches 64.52% in the combined model, which is equal to the sequence module and higher the regional module (45.16%). These results further confirm that the difference in the architecture of two attention mechanisms provides a complementary effect that can contribute to performance improvement.

**Figure 4.**
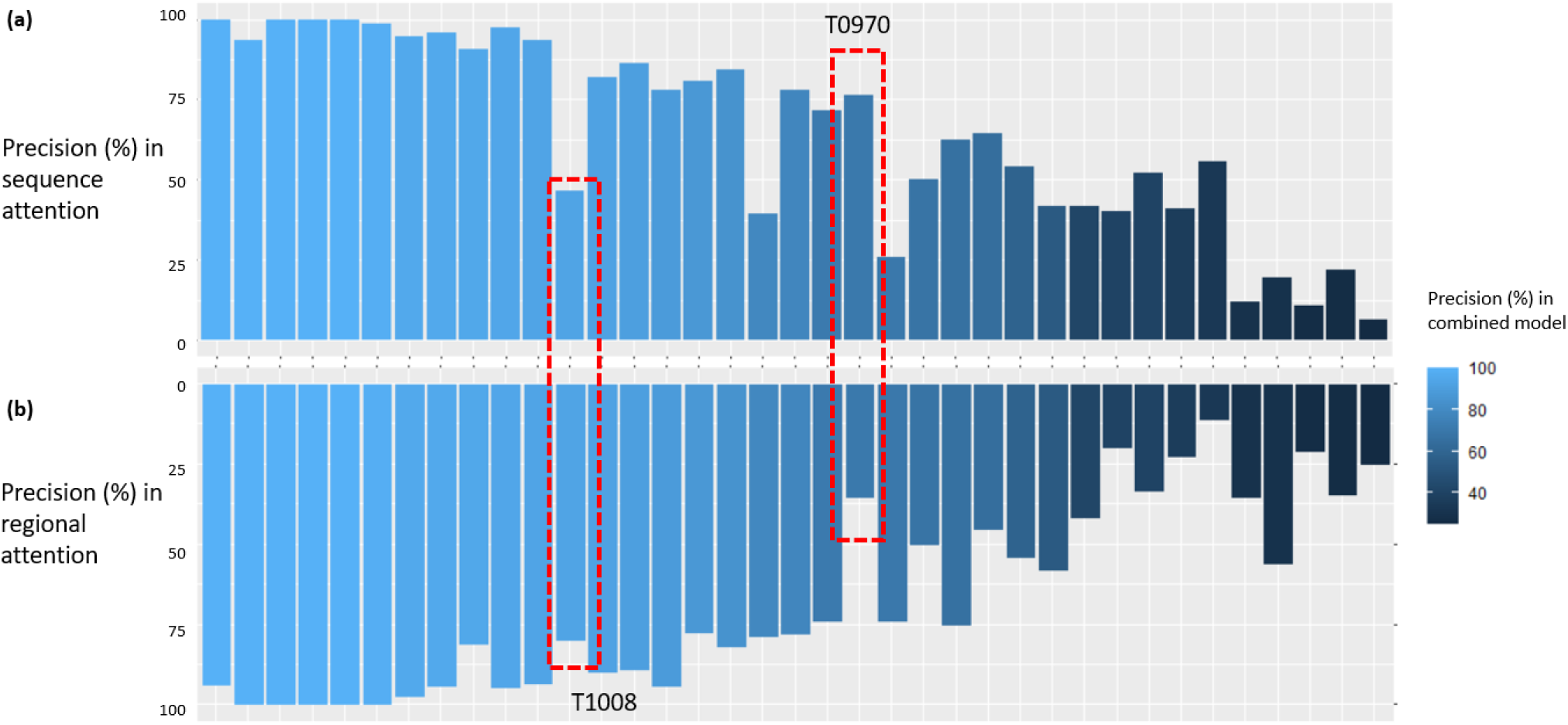
Comparison of the top-L/5 precision between sequence and regional attention module. The targets are arranged in the descending order of the top-L/5 precision in the combined model. **(a)** Precision scores from the sequence attention module. **(b)** Precision scores from the regional attention module. T1008 and T970 are two examples in which the two attention modules perform very differently.

To investigate the differences between the two attention modules, we select two targets: T1008 and T970, which have distinct performance patterns with the different attention implementations (**Table 2**). The top-L/5, L/2, and L precision of target T1008 in the regional attention module are all higher than the sequence attention module. The improvement of the performance suggests the contributions of more effective recognition of high-level 2D features from the regional attention mechanism. The regional attention module successfully identifies the contact pairs around the 26(V) and 65(E) regions (**Figure 5a**). In contrast, the sequence attention module assigned higher probability scores to a variety of false positive residue pairs in off-target areas (**Figure 5b**). Furthermore, we examine the attention weights from the top-L/2 predictions of the regional attention module. We add up the attention scores that are used to weight the same top-L/2 contacts for each prediction, resulting in an L×L map of normalized attention scores (**Figure 5c and 5d**). The result shows that the many of the contact pairs around the 26(V)-65(E) area have high weights in this map, which provides a possible explanation of why the predictions in the area are correct with the regional module.

**Figure 5.**
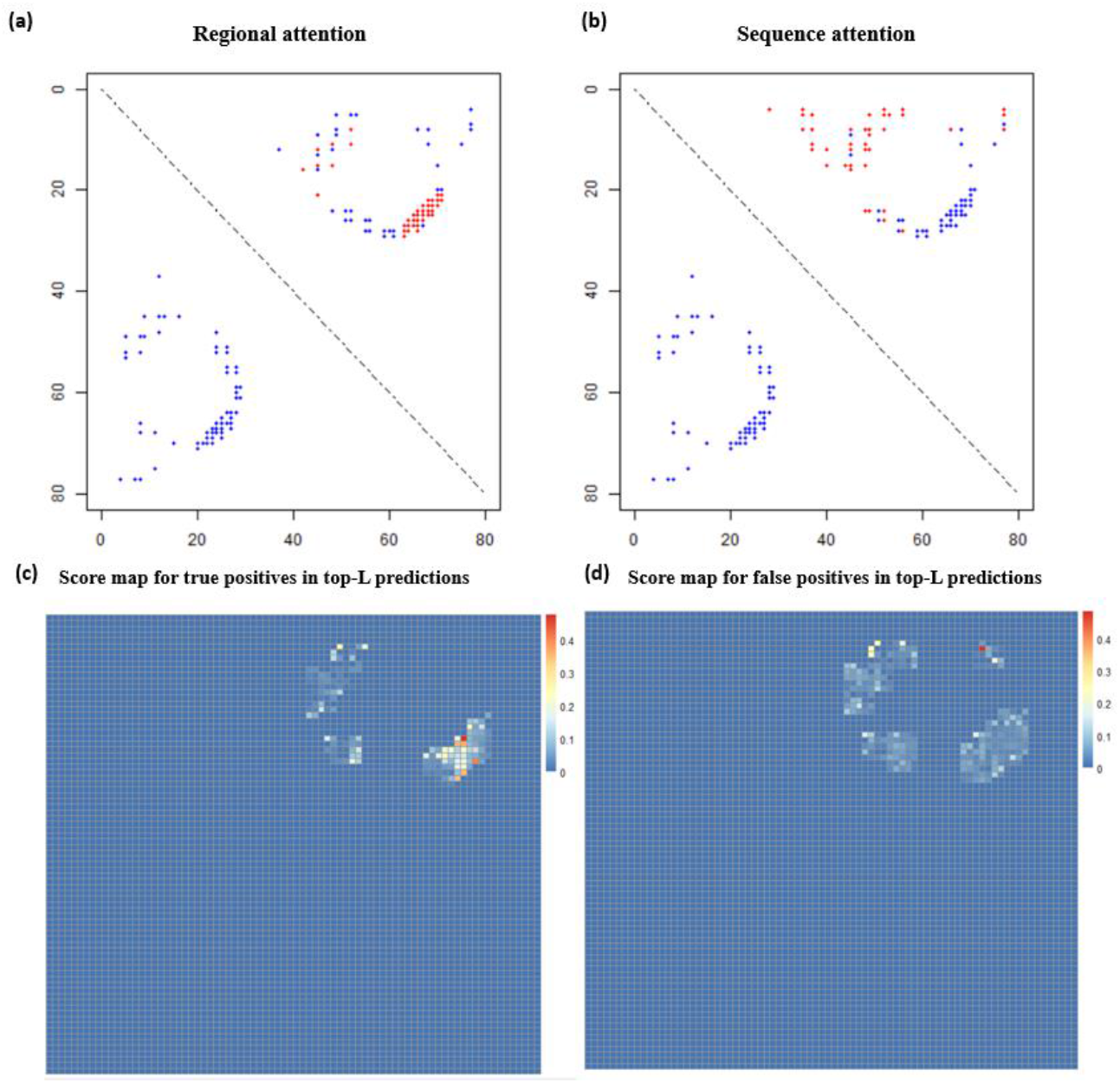
Comparison of performance between sequence and regional attention modules with long-range contacts of CASP13 target T1008. **(a)** Top-L/2 predictions from the regional attention module (red dots: contact predictions; blue dots: true contacts). **(b)** Top-L/2 predictions from the sequence attention module (red dots: contact predictions; blue dots: true contacts). **(c)** Visualization of attention scores from the true positives in top-L predictions. **(d)** Visualization of attention scores from the false positives in top-L predictions. For (a) and (b), contact predictions (red dots) are only shown in the upper triangle because of the symmetry of contacts, while true contacts are shown in both upper and lower triangle for better contrast. For (c) and (d), the sum of attention scores are normalized by the number of scores contributed to the sum of each position.

**Table 2.** Comparison of precision (%) on targets T1008 and T0970 between the two attention modules.

| Target | Model | Top-L/5 (%) | Top -L/2 (%) | Top -L (%) | Top -2L (%) |
| --- | --- | --- | --- | --- | --- |
| T1008 | Sequence | 46.67 | 43.59 | 25.97 | 15.58 |
|  | Regional | 80 | 64.1 | 42.86 | 24.68 |
| T0970 | Sequence | 76.47 | 72.09 | 56.47 | 35.29 |
|  | Regional | 35.29 | 39.53 | 35.29 | 25.88 |

Next, we examine target T970, which shows a better performance with the sequence attention module. The top-L/5, L/2, and L precision of T970 are 76.47%, 72.09%, and 56.47% in sequence attention module, respectively, much higher than the regional attention module (35.29%, 39.53%, and 35.29%, respectively). According to **Figure 6**, the sequence attention module correctly identifies the contact pairs between 5(F)-7(T) and 93(I)-95(F). In contrast, the regional attention module fails to recognize most of the area correctly. We use a similar technique to extract the 1D attention scores from the single attention module. For each residue, high sequence attention scores are observed where the true positive predictions are located (**Figure 7**), indicating the contributions of the 1D attention scores to the identification of these true positives.

**Figure 6.**
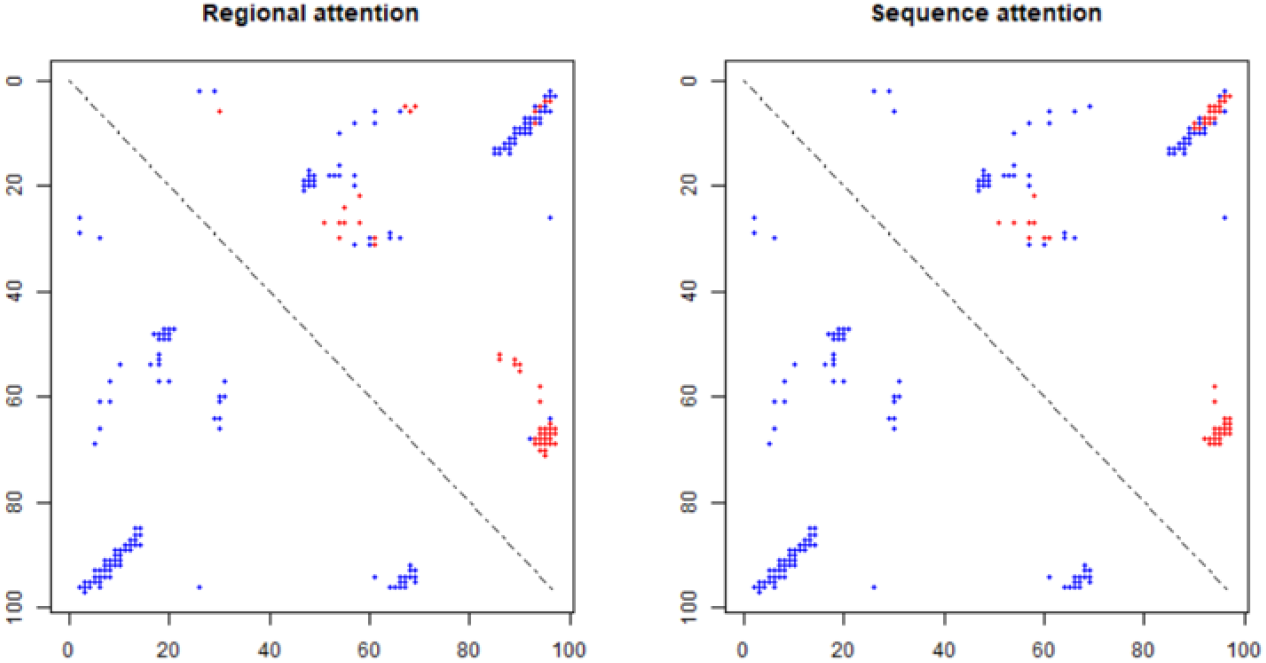
Comparison of performance between the sequence and regional attention module with long-range contacts for CASP13 target T970. The red and blue points indicate top-L/2 contact predictions and true contacts, respectively.

**Figure 7.**
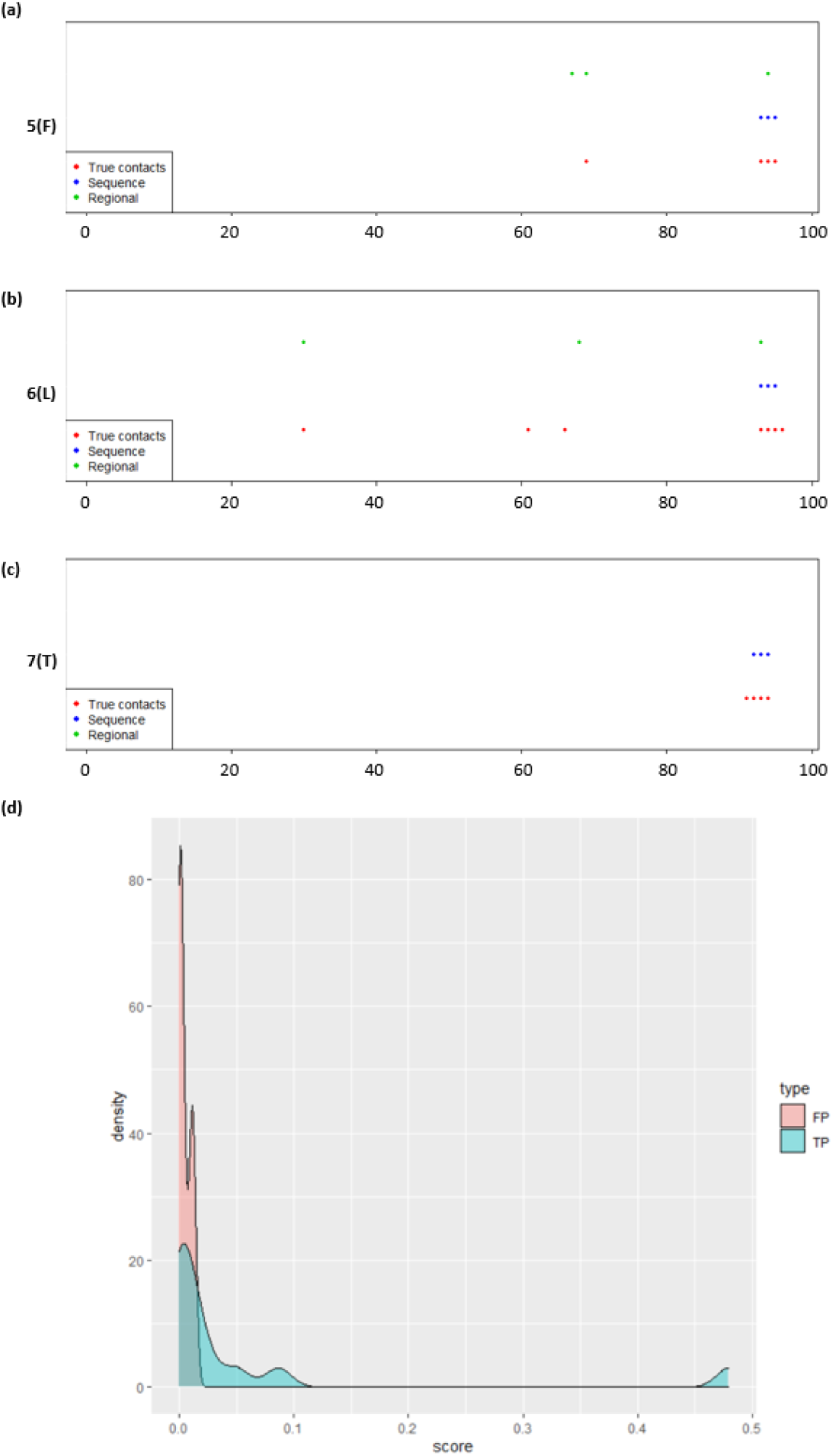
Visualization of top-L/2 predictions and corresponding true contacts of CASP13 target T970 from the sequence attention module. **(a), (b), (c)** illustrates true and predicted contacts for residue 5(F), 6(L), 7(T) along the protein sequence, respectively. Each point represents a true or predicted contact at the position of the other residue involving in the contact. **(d)** Distribution of the attention score of true positive and false positive contact predictions involving 5(F), 6(L), and 7(T). The attention scores are added up for all four heads at each position.

### 3.3 The effect of the regional attention score on contact prediction

To further investigate the influence of the intermediate attention layer on the regional attention mechanism, we extract the L×L feature map from the input of the attention layer in the regional attention module. Here we use two targets, T1005 and T0949, as examples, which have the highest top-L/5 scores in our predictions (**Figure 8**). We find that the primary effect of the attention mechanism is to reduce the values of false-positive contacts while increasing the scores for true positives for the input feature map. Furthermore, the comparison between the attention scores and predicted contact maps shows that the areas with higher attention scores are relatively sparser than contacts. Next, we consider the importance of the area with the high attention scores in contact prediction. To demonstrate this, we permute the input features around the positions that have high or low attention scores and use this permuted feature for prediction. Our results show that the number of true positive predictions will decrease most drastically (**Figure 9**), indicating that they contain important information related to protein fold. Also, the level of decrease remains similar when the region of permutated data grows from 1×1 to 5×5 in areas with high attention scores. In contrast, the level of decrease in areas with low attention score is much smaller and increases with the expansion of the permuted area. These results indicate the existence of potential protein folding-related key information in small areas with high attention scores.

**Figure 8.**
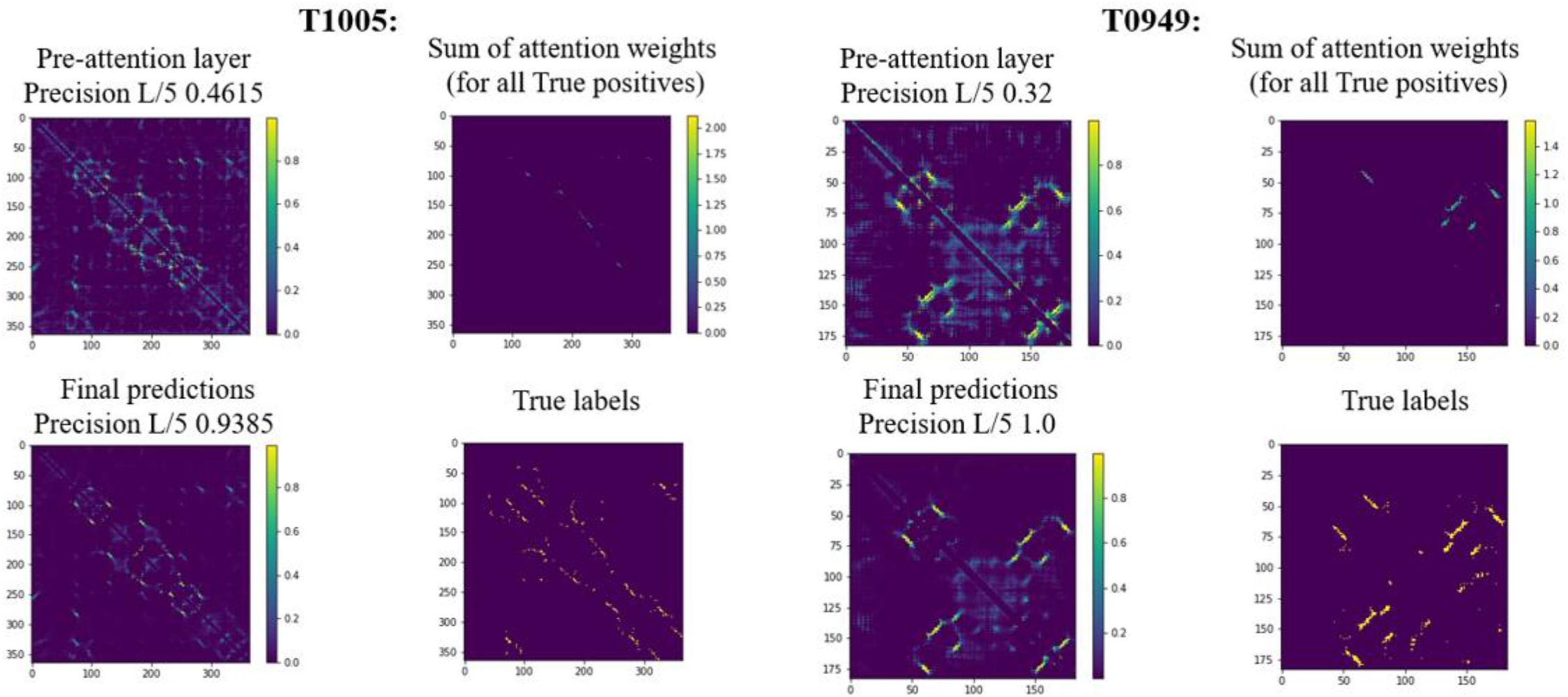
Visualization of data transformation of the attention layer of the regional attention module for CASP13 targets T1005 and T0949. For each target, we present the input feature map of the attention layer, attention weights, predicted scores, and true labels.

**Figure 9.**
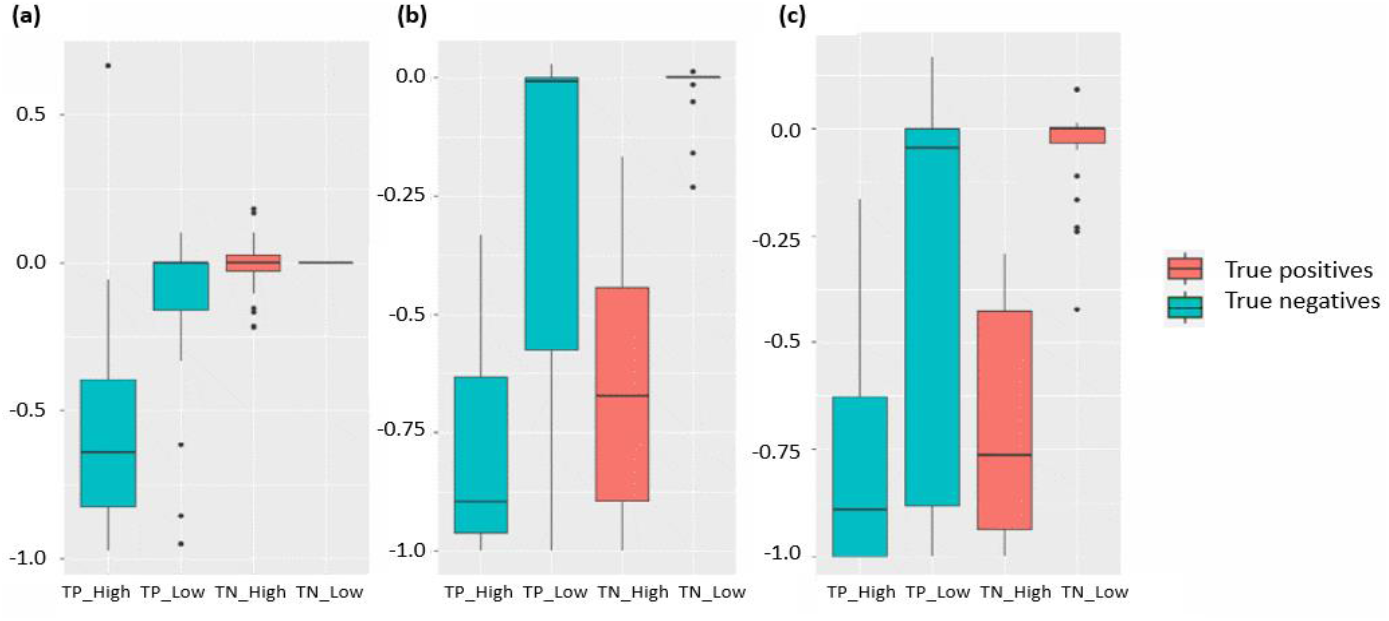
Performance after permutation of different locations of the input. The Y-axis indicates the increase or decrease of top-L/5 precision scores after permutation. Here we choose locations that have the highest and lowest k attention scores as centers for permutated regions, where k is the number of true positives of each target. **(a)** Impact of permutated regions with size (1,1). **(b)** Impact of permutated regions with size (3,3). **(c)** Impact of permutated regions with size (5,5). TP_High: true positive predictions with high scores. TP_Low: true positive predictions with low scores. TN_High: true negative predictions with high scores. TP_Low: true negative predictions with low scores.

### 3.4 Attention scores help to uncover the underlying key residues of protein folding

To further explore the interpretability of our method, we analyze the model on a protein whose folding mechanism has been well studied: Human common-type acylphosphatase (AcP). The structure and sequence information of AcP is obtained from PDB (https://www.rcsb.org/structure/2W4C). Vendruscolo et al. ^31^ identified three key residues in AcP (Y11, P54, and F94) that can form a critical contact network and result in the folding of a polypeptide chain to its unique native-state structure. The 3D structure model and three key residues are shown in **Figure 10a**.

**Figure 10.**
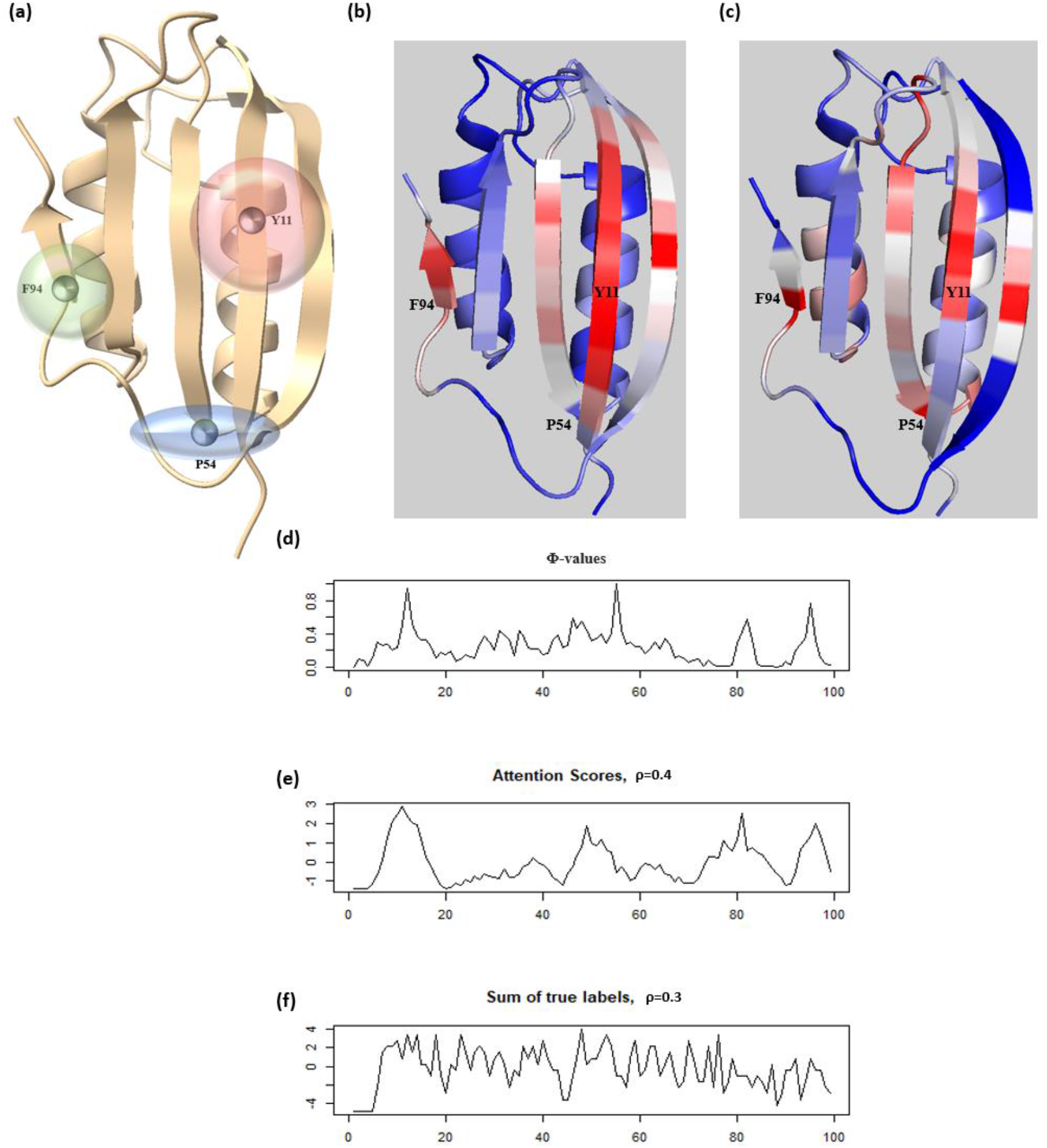
Visualization and interpretation contact predictions of Human common-type acylphosphatase from the regional attention module. **(a)** The 3D model of AcP with the three highlighted critical residues in protein folding. The transparent spheres around the residues indicate their corresponding scopes in the contact networks. **(b)** The heatmap of regional attention scores shown on the 3D structure of AcP. **(c)** The heatmap of Φ-values shown on the 3D structure of AcP. **(d), (e), (f)** The Φ-values, attention scores and the count of true contacts for each reside plotted along the protein sequence. ρ: Pearson Correlation Coefficient.

We use the regional attention module to predict the contact map of the protein. The precisions of the top-L/5, L/2, and L prediction are 100%, 95.74%, and 75.79%, respectively. We then extract the 2D attention score matrix from the model and combine the normalized row sums and column sums to reformat its dimension to L × 1. The attention score mapped to the protein 3D structure spot two key residues: Y11 and F94, where large regions of high attention weights are located (**Figure 10b**). Furthermore, we apply the same strategy with the experimentally determined Φ-values ^32^ (a value measuring the importance of the residue for protein folding) on the 3D structure of AcP (**Figure 10c)**. The Φ-values are the ratio of the change in stability of the transition-state ensemble (TSE) to that of the native state due to the mutation of each residue, and represents important information about residue interactions present within the TSE ^33^. The comparison (**Figure 10d, e**) shows that the Φ-values and normalized attention scores have similar trends along the peptide sequence (Pearson correlation coefficient = 0.4) with three peaks for Y11, P54, and F94 appeared in neighboring regions the curves determined by both the experimental method and the attention method. Also, we find that the true contract map does not provide the same level of information about the three key residues (**Figure 10f**). These results indicate that the attention scores can be applied to identify the critical components of the protein contact network important for protein folding when accurate contact predictions can be acquired. And this may also explain the importance of areas with high attention scores for successful contact prediction.

## 4 Conclusion

Interrogating the input-output relationships for complex deep neural networks is an important task in machine learning. It is usually infeasible to interpret the weights of a deep neural network directly due to their redundancy and complex nonlinear relationships encoded in the intermediate layers. In this study, we show how to use attention mechanisms to improve the interpretability of deep learning contact prediction models without compromising prediction accuracy. More interestingly, patterns relevant to key fold-determining residues can be extracted with the attention scores. These results suggest that the integration of attention mechanisms with existing deep learning contact predictors can provide a reliable and interpretable tool that can potentially bring more insights into the understanding of contact prediction and protein folding.

